# Transition metal activation reframes SAMHD1 regulation

**DOI:** 10.64898/2026.04.23.720456

**Authors:** Logan A. Calderone, Anthony Gizzi, Soumika Pinninti, James T. Stivers, Maria-Eirini Pandelia

## Abstract

SAMHD1 is the lone human dNTP triphosphohydrolase and is intimately linked to HIV viral restriction, dNTP pool maintenance, resistance to chemotherapy, and the autoinflammatory Aicardi-Goutières syndrome. While its substrate promiscuity and nucleotide basis of activity have been extensively studied, the identity and mechanistic roles of its metal cofactors remain poorly defined. Here, we integrate elemental analysis, spectroscopy, protein cross-linking, and enzyme kinetics to elucidate the molecular mechanisms underlying metal-dependent activation and catalysis in SAMHD1. Our findings establish that transition metals are essential components of SAMHD1 function, highlight their overlooked role in allosteric regulation, and reveal a central role for iron in organizing the dinuclear active site. We show that iron is preferentially incorporated in one position of the bimetallic core, where it promotes recruitment of a second divalent metal ion required for activity. While manganese can substitute for iron, it alters the metal binding equilibria, highlighting the unique functional properties of iron. Notably, SAMHD1 exhibits metal cofactor promiscuity at the second metal site, accommodating diverse divalent metals with distinct effects on activity. Cumulatively, our findings establish iron as a core structural and functional determinant of SAMHD1 catalysis and reveal how transition metal selectivity and flexibility enable enzymatic activity across diverse cellular environments and metal flux conditions.

**Significance Statement:** SAMHD1 is a central regulator of cellular dNTP poolsand an essential antiviral restriction factor, yet its metal dependence remains poorly defined. Here, we show that SAMHD1 is not a magnesium-driven enzyme but a transition-metal-dependent hydrolase in which iron plays a central structural and regulatory role. We define the metal requirements of the active and allosteric sites and demonstrate that diiron and heterodinuclear iron-containing cofactors form in solution and support catalysis. Transition metals such as iron and manganese act as more effective activators than magnesium, while plasticity at the second metal-binding site enables activity across dynamic metalation and oxidation states. SAMHD1 can thus flexibly tune antiviral defense and nucleotide metabolism, bypassing constraints imposed by metal availability and the cellular redox environment.

## Introduction

Metalloproteins account for more than 30% of all known proteins and exploit metal ions as structural elements, activators, and catalysts (1, 2). Despite their prevalence, metal cofactors are frequently misassigned or incompletely characterized, particularly in iron-containing proteins (3, 4). Owing to their redox activity, iron-containing metalloproteins are especially vulnerable to oxidation, cofactor degradation, and incomplete metal occupancy, further complicating their functional annotation (3, 5, 6). Historically, iron has been primarily associated with oxidases and oxygenases, in which its redox versatility is harnessed to activate molecular oxygen and enable chemically challenging transformations (3, 7, 8).

More recently, iron has emerged as a functionally important cofactor in non-redox enzymes such as hydrolases (3, 9). These findings have challenged the hitherto accepted notion that iron is incompatible with redox-inert reactions and instead suggest that it may be selectively employed to support catalysis different cellular niches and metal availabilities. Consistent with this view, a growing number of phosphatases and phosphodiesterases, including purple acid phosphatases, calcineurins, Cas3 nucleases, and cyclic dinucleotide hydrolases, have been shown to exhibit iron-dependent activity (10–13). This trend is particularly pronounced within the HD-domain superfamily, which encompasses both oxidative and hydrolytic enzymes (9, 14). An increasing number of hydrolases within this family have been shown to coordinate iron, potentially reflecting an evolutionary legacy of mixed chemistries within this protein fold (3, 12, 13, 15).

One such HD-domain hydrolase in which iron has been identified as a constituent of the active site, but in whichthe identity and role of metals in activity remains poorly defined, is the sterile alpha motif and HD-domain containing protein 1 (SAMHD1) (16–21). SAMHD1 is the only human deoxyribonucleotide triphosphate (dNTP) (metallo)triphosphohydrolase and is intimately involved in restriction of viruses, including HIV; dNTP pool maintenance, resistance to chemotherapy, as well as the autoinflammatory Aicardi-Goutières syndrome (17, 18, 22–32). These diverse functions depend on metal-dependent oligomerization mediated by two allosteric sites, one (d)GTP specific and the other dNTP dependent, as well as metal-dependent catalysis at the active site (32–36). While the structural determinants and functional consequences of oligomerization have been extensively studied, the identity, occupancy, and mechanistic role of the active and allosteric site metals remain incompletely defined. As a result, how metal coordination governs SAMHD1 assembly and enzymatic function remains poorly understood.

Here, we integrate biochemical assays with metal-specific spectroscopic analyses of selectively metalated SAMHD1 forms to define the metal requirements underlying its dNTP phosphohydrolase activity. Using metal-depleted (apo) SAMHD1, we establish that catalytic activity strictly requires a transition metal ion. Reconstitution experiments further reveal that SAMHD1 readily incorporates iron, assembling either redox-active diiron or heterodinuclear active sites with other divalent metals. Within the bimetallic active site core one metal-binding site exhibits strong selectivity for iron, whereas the second site is more permissive, accommodating various divalent ions with functional consequences ranging from activation to inhibition. This metal tolerance extends to the allosteric sites, in which iron and manganese activate SAMHD1 with higher affinity than the canonical magnesium activator (23, 24, 33–38).

Focusing on the active site, we show that the iron oxidation state does not greatly influence activity in heterodinuclear Fe-M^2+^ assemblies but controls catalysis in the homodinuclear Fe-Fe configurations. Moreover, iron uniquely promotes cooperative assembly of a dinuclear center, a property not recapitulated by manganese. Thus, iron acts as a critical determinant of both active-site architecture and function. Collectively, these findings place SAMHD1 within a growing class of iron-dependent hydrolases and underscore an underappreciated role for redox-active metals in non-redox enzymatic reactions. Our results prompt a reassessment of the canonical metallocofactor assigned to SAMHD1 and suggest that iron may confer functional advantages in regulating dNTP catabolism. Notably, the strong specificity and enhanced activation by iron now establishes that dNTP degradation by SAMHD1 is iron dependent, similar to dNTP biosynthesis by ribonucleotide reductase (39–41).This work advances our understanding of metal-regulated nucleotide metabolism and opens new avenues for therapeutic intervention in immune regulation and cancer (26, 27, 38).

## Results

### SAMHD1 catalysis requires transition metal ions

To determine the metal dependence of SAMHD1 activity, we generated a metal-depleted form of SAMHD1 (apo-SAMHD1) and assessed dGTP turnover in the presence of Mg^2+^, Fe^2+^, or Mn^2+^ (**Fig.1A**). In line with previous studies, no dNTP hydrolysis activity was detected in the absence of metal ions. Although Mg^2+^ is considered the canonical activator, its addition only resulted in a slow hydrolysis rate with minimal activity enhancement upon increasing its concentration (23, 24, 33–38). In contrast, Fe^2+^ and Mn^2+^ activated SAMHD1 with higher affinity and greater reaction velocities, demonstrating that these transition metals activate the enzyme more effectively (**Fig. 1A**). Apo-SAMHD1 contained trace amounts of Fe^2+^ (**Table S1**), suggesting that the low activity observed upon Mg^2+^ supplementation likely arises from residual iron contamination.

**Figure 1.**
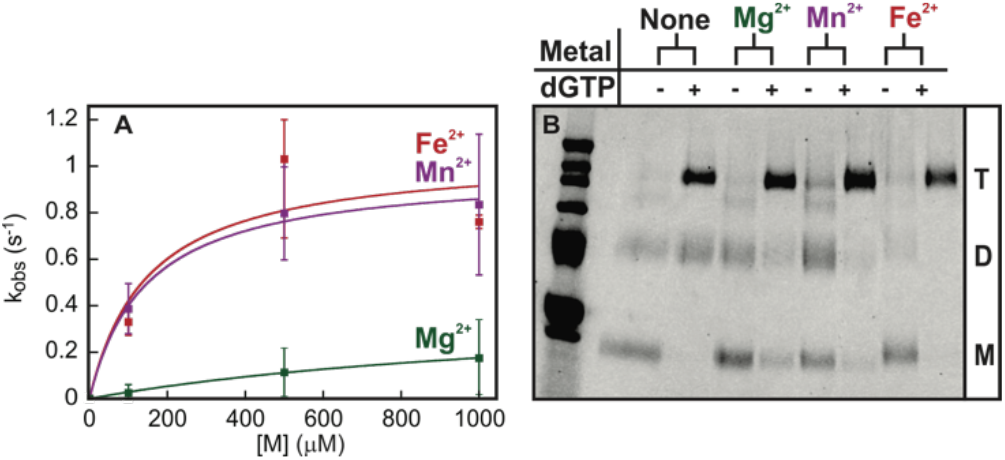
Metal dependence of oligomerization and activity in SAMHD1. **A**. Dependence of apo-SAMHD1 activity on concentration of Fe^2+^ (red, activation constant *K*_act_ = 150 ± 168 µM), Mn^2+^ (purple, *K*_act_ = 146 ± 30 µM), or Mg^2+^ (green, *K*_act_ = 1350 ± 200 µM). The experiment was performed anaerobically under reducing conditions. **B**. Glutaraldehyde crosslinking gel depicting oligomerization of SAMHD1 by Mg^2+^, Mn^2+^, or Fe^2+^, with and without dGTP. The T, D, and M labels indicate tetramer, dimer, and monomer respectively. Error bars indicate the standard error of the mean (n = 2).

To assess the relationship between metal binding and oligomerization, we employed glutaraldehyde chemical cross-linking to confirm the SAMHD1 oligomeric state in the activity assays (33, 42). In the absence of added metal ions, dGTP alone promoted partial tetramer formation, whereas addition of divalent metals further enhanced oligomerization (**Fig. 1B, Fig. S1**). Because the dGTP stock was essentially metal free (**Table S1)**, these observations indicate that nucleotide binding can drive a basal level of tetramerization independent of an allosteric metal, hitherto proposed to be Mg^2+^ coordinating the two nucleotides (23, 24, 33–38). However, full conversion to the tetrameric form required the presence of a metal ion, consistent with a concerted mechanism in which both nucleotide binding and metal coordination cooperate to stabilize the active oligomer (23, 34). Both Fe^2+^ and Mn^2+^ markedly enhanced SAMHD1 catalytic activity and promoted formation of the tetramer. In contrast, Mg^2+^ supported stable tetramer assembly but resulted in only minimal catalytic turnover **(Fig. 1A-B)**. The data demonstrate that oligomerization alone is not sufficient for full enzymatic activation. Rather, robust catalysis requires iron or manganese. The limited activity observed in the presence of Mg^2+^ is consistent with catalysis arising from mixed-metal or incompletely occupied active sites, as suggested by recent crystal structures and elemental analyses (**Table S1**) (16, 17, 21, 23, 34).

### SAMHD1 assembles diverse Fe-X dinuclear active sites

Our experiments with apo-SAMHD1 combined with recent crystallographic data suggest that SAMHD1 must be capable of assembling different dinuclear cofactors. To directly probe formation of such sites, we turned to EPR spectroscopy. Selective enrichment of SAMHD1 with iron was achieved by heterologous expression in minimal media supplemented with iron (M9-Fe) followed by isolation under O_2_-free conditions (**Table S1)**. The samples were treated with sodium ascorbate to probe the accumulation of any mixed-valent paramagnetic states (43–45). Indeed, the spectrum of the ascorbate-reduced M9-Fe SAMHD1 showed signals consistent with the formation of antiferromagnetically-coupled (*S* = 1/2, g_av_ < 2) mixed-valence (Fe^3+^-Fe^2+^) diiron centers (**Fig. 2A**) (43–47). A minor signal at g = 4.3 was also detected, corresponding to a high-spin mononuclear Fe^3+^ species accounting for less than 5% of the total iron content (**Fig. 2B**) (48).

**Figure 2.**
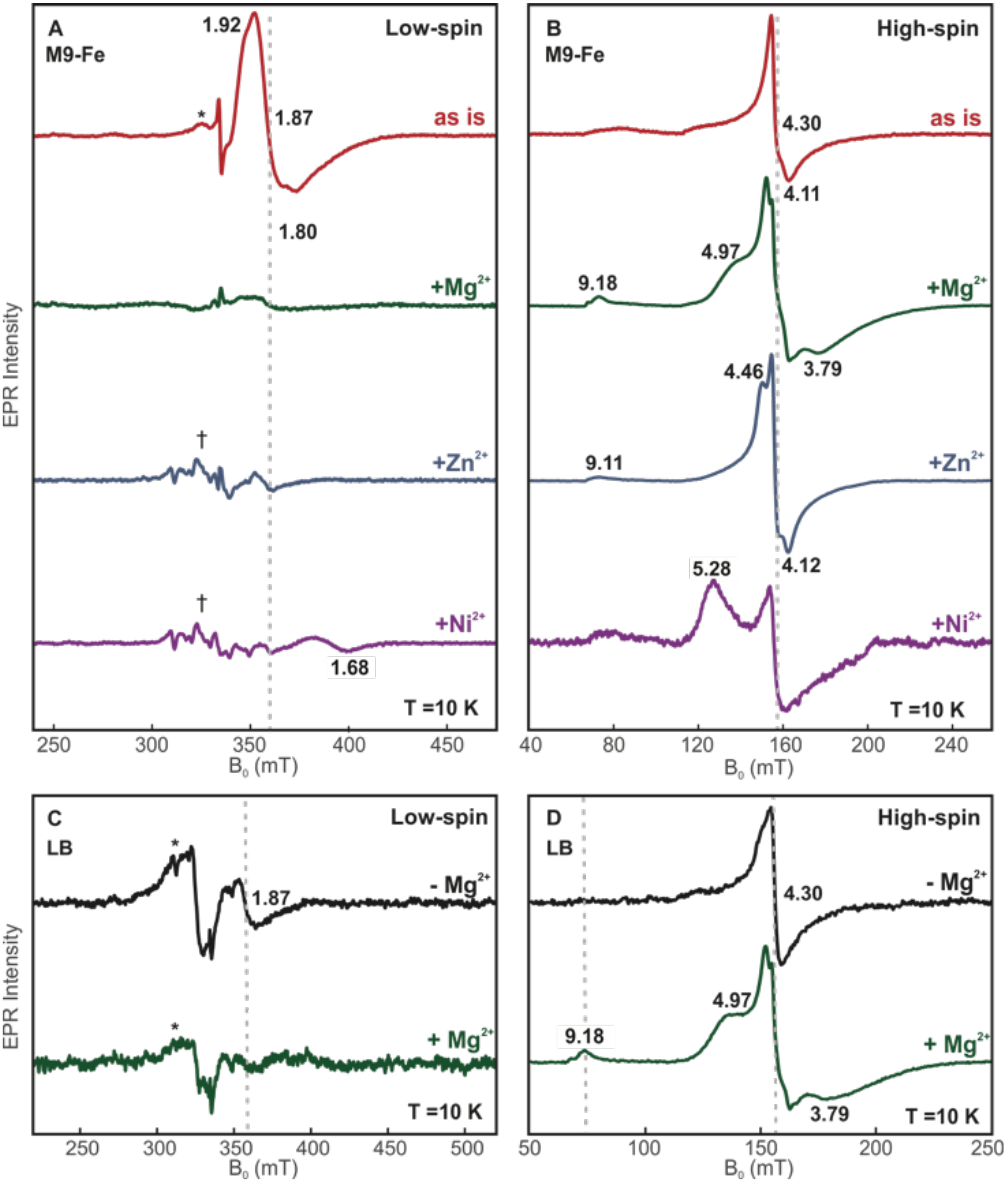
Monitoring assembly of a diiron cofactor and incorporation of various divalent metal ions at the active site of SAMHD1 by EPR. **A**. EPR spectra of M9-Fe SAMHD1 as is (red) or with supplemental Mg^2+^ (green), Zn^2+^ (blue), or Ni^2+^ (purple) in the low-spin region. **B**. EPR spectra of the same samples as in panel **A** in the high-spin region. **C**. EPR spectra in the low-spin region of LB-SAMHD1 purified in either the absence (black) or presence (green) of Mg^2+^. **D**. EPR spectra of the same samples in panel **C** in the high-spin region. Asterisks indicate signals from Cu^2+^ contamination. Daggers indicate signals from Mn^2+^ contamination. Experimental conditions: temperature = 10 K, microwave frequency = 9.38 GHz, microwave power = 2 mW, modulation amplitude = 1 mT.

Addition of divalent metal ions markedly altered the EPR spectra in both the low-spin and high-spin regions. Supplementation with Mg^2+^, Zn^2+^, or Ni^2+^ led to the loss of the g_av_ < 2 diiron signal and the appearance of new high-spin resonances near the mononuclear Fe^3+^ region **(Fig. 2A-B)**. Notably, these resonances exhibited pronounced rhombicity distinct from that of the isolated mononuclear Fe^3+^ species (**Fig. S2A-B**). The spectral changes are consistent with displacement of diiron centers and formation of mixed-metal, iron-containing dinuclear sites (49–51). Specifically, addition of Mg^2+^ produced signals at g = 9.18, 4.97, and 3.79 arising from excited and ground state Kramers’ doublets, which are consistent with changes in the rhombicity of the high-spin Fe^3+^ due to its complexation with a divalent metal (**Fig. 2B, Fig. S2B**). Addition of Zn^2+^ resulted in the appearance of a high-spin feature at g ~ 9.11 as well as additional resonances in the region of the mononuclear Fe^3+^ signal, while addition of Ni^2+^ resulted in a rhombic signal with apparent g values of 5.28 and 1.68 (**Fig. 2B)**. Together, these data demonstrate that SAMHD1 readily remodels its dinuclear active site in response to metal availability, forming multiple iron-based dinuclear cofactors with a variety of metal ions.

To reconcile these findings with earlier studies in which metal supplementation was not specified or controlled, we next analyzed SAMHD1 purified from LB media by EPR spectroscopy with and without exogenous Mg^2+^ added during purification. When Mg^2+^ was omitted, the resulting spectra displayed a low intensity g_av_ < 2 signal, consistent with formation of diiron centers (**Fig. 2C**). In contrast, inclusion of Mg^2+^ in the purification buffers, as employed in prior protocols (17, 23, 24, 52), abolished this mixed-valence Fe^3+^-Fe^2+^ signal, indicating suppression of diiron site assembly. Notably, the high-spin signal of the Mg^2+^-supplemented LB-derived SAMHD1 closely matched that observed for Mg^2+^-treated M9-Fe samples, further confirming its assignment to Fe^3+^-Mg^2+^ mixed-metal centers (**Fig. 2D**). The observed spectroscopic signatures directly correspond to metal configurations previously inferred from crystallographic analyses, providing the first experimental evidence that the dinuclear metal sites reported in crystalsare populated in solution and are modulated by metal availability during purification (17, 18, 21).

### Manganese-dependent cofactor assembly and catalytic activation

Having established that iron directs the assembly of multiple dinuclear cofactors in SAMHD1, we next examined how perturbation of the metal-binding environment influences cofactor integrity. Guided by crystallographic models identifying His233 as a ligand of the more permissive metal site, we used EPR spectroscopy to assess the impact of removing this residue (17, 18, 21). Substitution of H233 with alanine abolished formation of the dinuclear cofactor, confirming its essential role in metal coordination and cofactor assembly (**Fig. 3A, 3B**).

**Figure 3.**
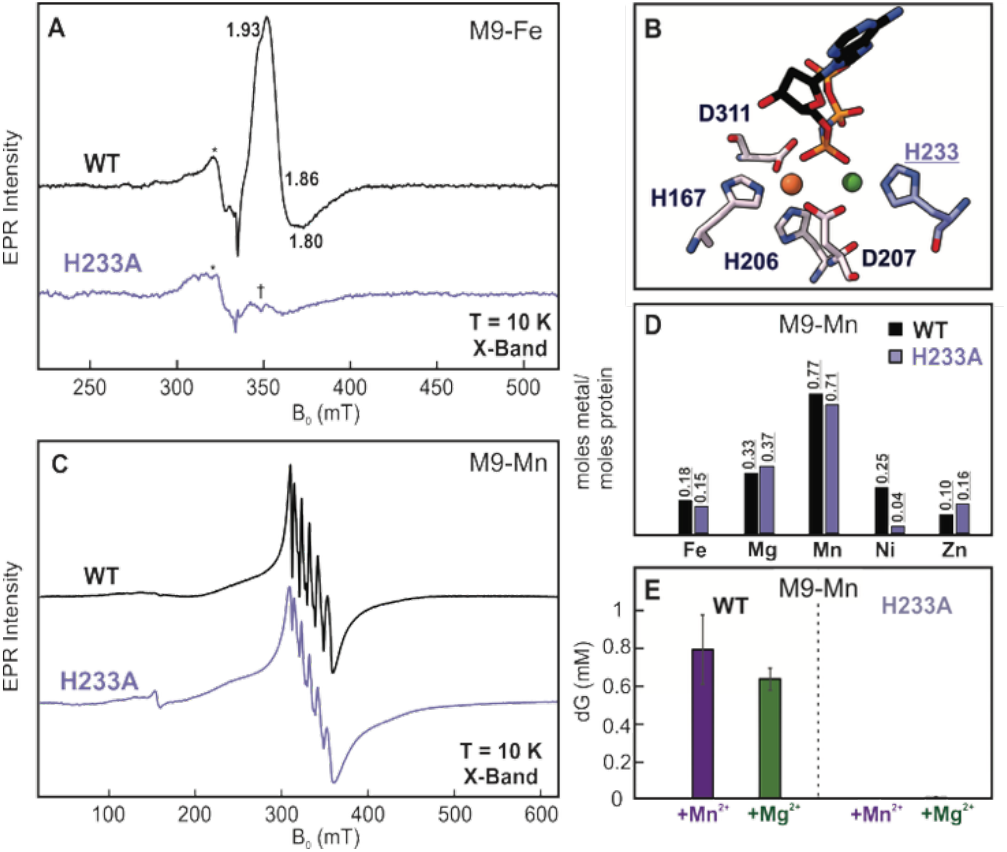
Spectroscopic characterization of wild-type and H233A SAMHD1 under different metalation enrichment, metal content and catalytic activity of Mn-enriched SAMHD1. **A**. EPR spectra of wild-type and H233A SAMHD1 enriched with iron (M9-Fe). Asterisks indicate signals from Cu^2+^ contamination. Daggers indicate signals from Mn^2+^ contamination. **B**. Active site structure of the *Hs* SAMHD1 with 2’-deoxy-5’-O-[(R)-hydroxy{[(R)-hydroxy(phosphonooxy)phosphoryl]amino}phosphoryl] adenosine bound (PDB ID: 6TX0). The Fe ion is shown as an orange sphere and the Mg ion as a green sphere. Only the primary coordinating ligands are shown with the H233 ligand to the Mg ion (site 2) highlighted in purple. **C**. EPR spectra of wild-type and H233A SAMHD1 enriched with manganese (M9-Mn). **D**. ICP-AES data comparing the molar ratios of metals that copurify with wild-type (black bars) and H233A M9-Mn SAMHD1 (purple bars). **E**. HPLC endpoint activity assays comparing the activity of wild-type and H233A M9-Mn SAMHD1 with supplemental Mn^2+^ (purple bars) or Mg^2+^ (green bars). Experimental conditions: temperature = 10 K, microwave frequency = 9.38 GHz, microwave power = 2 mW, modulation amplitude = 1 mT. Activity assays were carried out with [*Hs* SAMHD1] = 0.5 μM, [dGTP] = 1 mM, and [Mn] = 2 mM or [Mg] = 2.5 mM for 10 min. Error bars indicate the standard error of the mean (n = 4).

To examine the effect of manganese on cofactor composition, wild-type and H233A SAMHD1 was selectively enriched with Mn^2+^ via heterologous expression in minimal media (M9-Mn, **Table S1**). EPR analysis revealed spectra dominated by a well-resolved six-line hyperfine pattern characteristic of mononuclear Mn^2+^ centers, rather than the broadened multiline features expected for dimanganese sites (**Fig. 3C**) (53, 54). This spectral signature was reproducible across multiple preparations, indicating that formation of a dimanganese cofactor is disfavored under these conditions and suggesting that Mn^2+^ occupancy at one site influences metal binding at the second one. The Mn^2+^-enriched H233A variant exhibited EPR spectra comparable to those of the wild-type enzyme, and ICP-AES analysis confirmed similar overall manganese content in both proteins (**Fig. 3D, Table S1**). Together, these data support a model in which manganese preferentially occupies the 4-coordinate HD site, whereas the H233-associated site displays reduced affinity for Mn^2+^ (**Fig. 3B)** (17, 18, 21).

Consistent with our previous observation that apo-SAMHD1 supplemented with Mn^2+^ is catalytically active (**Fig. 1A**), addition of Mn^2+^ or Mg^2+^ to wild-type M9-Mn SAMHD1 restores activity to rates comparable to those reported previously (0.5-5.0 s^−1^) (**Fig. 3E, Table S2)** (17, 33, 55). Although Mn^2+^-enriched SAMHD1 purified under these conditions predominantly harbors mononuclear Mn^2+^ centers, inclusion of excess Mn^2+^ (or Mg^2+^) during the activity assays promotes formation of catalytically competent dimetal sites. In contrast, the H233A M9-Mn variant remained inactive with Mn^2+^ (**Fig. 3E, Table S2)**, indicating that catalytic activity depends on assembly of a dinuclear cofactor and cannot be supported when coordination at the second metal-binding site is disrupted.

### Active-site iron and transition metals enhance SAMHD1 catalysis

To evaluate the impact of active-site metal identity on SAMHD1-catalyzed dNTP hydrolysis, we systematically varied the second metal in the dinuclear active center while keeping the allosteric effector constant (**Fig. 4**). We examined Fe^2+^-M^2+^ active-site configurations, where M was Mg^2+^, Mn^2+^, or Fe^2+^, using Mg^2+^ as the allosteric effector. All samples were pre-reduced with sodium dithionite to ensure that iron remained in the ferrous state for consistency. EPR spectroscopy confirmed that the excess Mg^2+^ added as effector did not exchange into the active site or alter its metal composition over the course of the assay (**Fig. S3, Fig. S4**). Because metal incorporation differed among preparations, catalytic rates were normalized to the iron content of each sample. Under these conditions, the Fe^2+^-Mg^2+^ form hydrolyzed dGTP with an activation constant (*K*_act_) of 257 ± 80 µM and an apparent rate of 1.8 ± 0.2 s^−1^, comparable to previously reported measurements (**Fig. 4A, Table S3**) (17, 33, 55). The homodinuclear Fe^2+^-Fe^2+^ cofactor supported indistinguishable catalytic activity and a similar activation constant, indicating that this configuration is fully competent for catalysis. In contrast, the Fe^2+^-Mn^2+^ form displayed a comparable *K*_act_ but slower catalytic turnover, suggesting a lower turnover efficiency under these conditions (**Fig. 4A, Table S3**). It should be noted that this preparation contained higher levels of Ni^2+^ than the others, which may partially suppress the observed rates (**Table S1**). Together, these results show that substitution at the second metal site does not substantially affect activation, and maximal catalytic rates are achieved with Fe^2+^-Fe^2+^ and Fe^2+^-Mg^2+^ active-site configurations.

**Figure 4.**
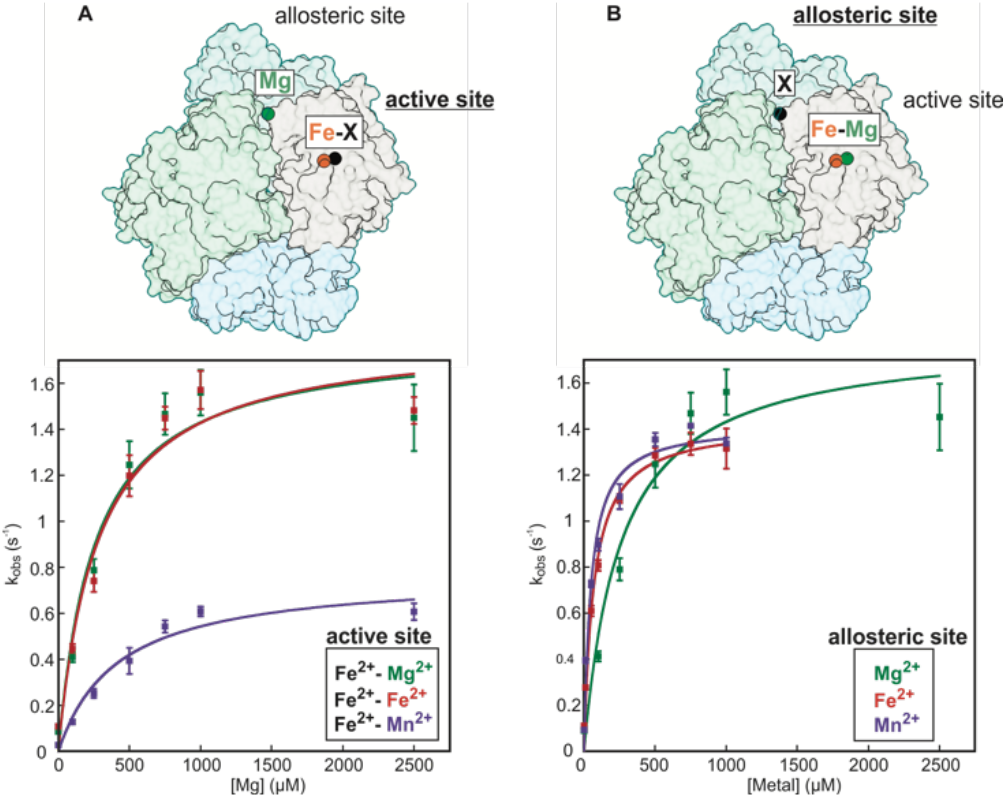
SAMHD1 activity assays highlight metal plasticity at the active and allosteric sites. **A**. HPLC activity assay of SAMHD1 enriched with Fe^2+^-Mg^2+^ (M9-FeMg, green, k_obs_ = 1.80 ± 0.2 s^−1^, K_act_ = 257 ± 80 µM), Fe^2+^-Fe^2+^ (M9-Fe, red, k_obs_ = 1.82 ± 0.2 s^−1^, K_act_ = 277 ± 80 µM), or Fe^2+^-Mn^2+^ (M9-FeMn, purple, k_obs_ = 0.77 ± 0.08 s^−1^, K_act_ = 417 ± 120 µM) using magnesium as an allosteric activator. **B**. HPLC activity assay of SAMHD1 enriched with Fe^2+^-Mg^2+^ (M9-FeMg) using Mg^2+^ (green, k_obs_ = 1.80 ± 0.2 s^−1^, K_act_ = 257 ± 80 µM), Fe^2+^ (red, k_obs_ = 1.42 ± 0.05 s^−1^, K_act_ = 68 ± 10 µM), or Mn^2+^ (purple, k_obs_ = 1.42 ± 0.07 s^−1^, K_act_ = 47 ± 10 µM) as allosteric activators. Error bars indicate the standard error of the mean (n = 3).

We next examined the contribution of the allosteric metal effector to catalysis. Using the Fe^2+^-Mg^2+^ active-site configuration, dGTP hydrolysis was measured while varying the allosteric activator between Mg^2+^, Fe^2+^, and Mn^2+^ (**Fig. 4B**). Activation with Fe^2+^ supported a comparable maximal rate (*k*_cat_ = 1.42 ± 0.05 s^−1^) but with an approximately fourfold lower *K*_act_ (68 ± 10 μM) compared to Mg^2+^ (*K*_act_ = 257 ± 80 μM), indicating tighter allosteric activation. Furthermore, Mn^2+^ produced nearly identical kinetic parameters to Fe^2+^ (**Table S3**), demonstrating that transition metals can function as higher-affinity allosteric activators than Mg^2+^ and may therefore be physiologically relevant under cellular conditions. Together, the data reveal substantial metal-dependent flexibility in SAMHD1 catalysis, with various iron-containing active sites showing similarly robust turnover and transition metals supporting enhanced allosteric activation relative to Mg^2+^, challenging the notion of Mg^2+^ as the exclusive physiological allosteric activator.

### Role of iron oxidation state in SAMHD1 catalysis

The requirement for iron in catalysis prompted us to examine whether its redox state modulates SAMHD1 activity. This question is particularly relevant because SAMHD1 is expressed in macrophages (27, 56–60), a comparatively oxidizing cellular environment, and oxidation to the ferric state is inhibitory in some hydrolases (3). To define the accessible redox states of the SAMHD1 diiron cofactor, we employed ^57^Fe Mössbauer spectroscopy (**Fig. 5A, Table S4**). The Mössbauer spectra revealed that anaerobically purified SAMHD1 contained a mixture of oxidation states, consisting predominantly of the diferric (Fe^3+^-Fe^3+^, ~50%) and mixed-valent (Fe^3+^-Fe^2+^, ~30%) species (**Table S4)**. The remaining spectral intensity could not be completely accounted for due the broad background. Upon reduction with ascorbate, the mixed-valent Fe^3+^-Fe^2+^ state became the dominant species (~56%), consistent with the EPR measurements described above (**Fig. 2**). In this sample, a minor fraction of the diferric state (~10%) persisted, while the fully reduced diferrous (Fe^2+^-Fe^2+^) form accounted for ~30% of the iron population (**Table S4)**. Treatment with sodium dithionite resulted in quantitative conversion to the diferrous state. Thus, the SAMHD1 diiron cofactor populates diferric, mixed-valent, and diferrous states that can be interconverted by mild redox treatments (**Fig 5A, Table S4)**. This behavior closely parallels that reported for other HD-domain diiron enzymes and provides a framework to assess how the iron oxidation state influences catalytic function (12, 13, 15, 43, 44, 61).

**Figure 5.**
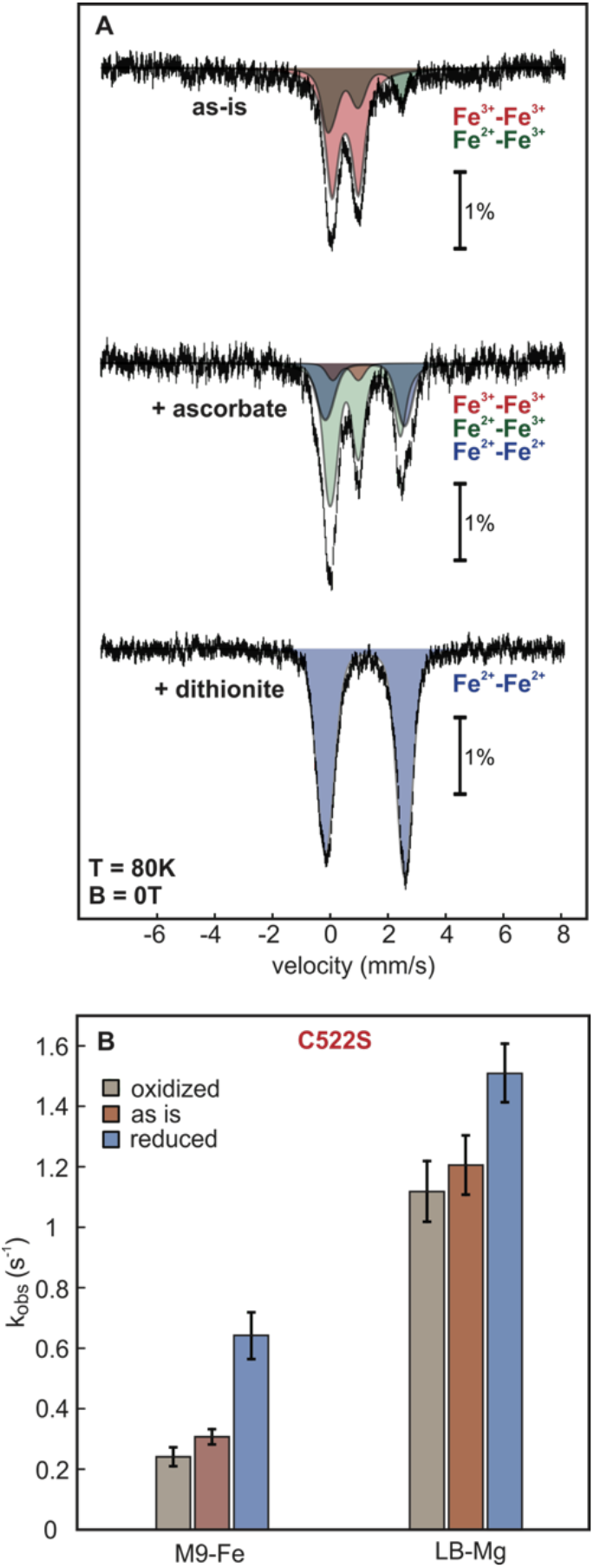
Distribution of redox states in Fe-enriched SAMHD1 by Mössbauer spectroscopy and redox-dependence of hydrolysis. **A**. Mössbauer spectra of M9-Fe SAMHD1 under various redox conditions showing individual spectra of the Fe^3+^-Fe^3+^ species (red), the Fe^3+^-Fe^2+^ species (green), and the Fe^2+^-Fe^2+^ species (blue). Overlapping spectra are shown as the convolution of the underlying colors. Raw data is displayed as black vertical bars. **B**. HPLC activity assay data of M9-Fe C522S SAMHD1 and LB-Mg C522S SAMHD1 under oxidizing (gray), as is (brown), or reducing conditions (blue). Error bars indicate the standard error of the mean (M9-Fe oxidized, as is, n = 5), (M9-Fe reduced, n = 6). (LB-Mg oxidized, as is, n = 7), (LB-Mg reduced, n = 8).

To further explore the effects of the Fe redox state on the overall catalytic activity of SAMHD1 we employed M9-Fe C522S SAMHD1 **(Table S1)** (**Fig. 5B)**. The C522S variant bypasses the known inhibition of the wild-type enzyme under oxidative conditions due to disulfide formation involving C522, while retaining wild-type activity (62–64). The as isolated protein hydrolyzed dGTP with an apparent rate of 0.31 ± 0.03 s^−1^, while treatment with hydrogen peroxide resulted in a slight decrease of the hydrolysis rate (**Fig. 5B, Table S5**). This persistent background activity even in an oxidizing environment likely stems from residual Mg^2+^ contamination that could not be fully eliminated (**Table S1**). In contrast, reduction with sodium dithionite, which produces a homogeneous diferrous center, resulted in an approximately two-fold increase in the hydrolysis rate (**Table S5)**.

We next asked whether a similar redox dependence is observed for heterodinuclear active sites. To this end, the Fe-Mg-enriched C522S SAMHD1 variant was examined under the same redox conditions. Oxidation with hydrogen peroxide, which maximizes the Fe^3+^-Mg^2+^ configuration, supported dGTP hydrolysis with an apparent rate of 1.12 ± 0.1 s^−1^, only slightly lower than that of the as-isolated enzyme (**Table S5)**. Further, reduction with sodium dithionite led to a modest increase in activity. Thus, in contrast to the diiron form, catalytic activity of the Fe-Mg^2+^ center is largely insensitive to the oxidation state.

Together, these data reveal that the susceptibility of SAMHD1 to redox-dependent inhibition depends strongly on the identity of the second metal ion. Diiron active sites exhibit pronounced redox sensitivity, with maximal activity in the diferrous state, whereas mixed-metal Fe-Mg^2+^ centers retain robust activity even under oxidizing conditions. These observations also imply that the mixed-valent Fe^3+^-Fe^2+^ state remains catalytically competent, consistent with our EPR experiments showing substrate binding to both the Fe^3+^-Fe^2+^ and Fe^3+^-Mg^2+^ forms (**Fig. S4, Fig. S5**) More broadly, incorporation of a redox-inert metal at the second coordination site buffers the enzyme against oxidative inactivation, providing a potential mechanism by which SAMHD1 maintains activity across different cellular redox environments.

## Discussion

Despite extensive investigation of SAMHD1 regulation and substrate specificity, the chemical identity and functional role of its metal cofactors have remained unresolved (16–21, 65, 66). Structural studies suggested a dinuclear Fe-Mg^2+^ center within the catalytic HD domain, yet the contributions of individual metals, their oxidation states, and roles in catalysis have not been resolved (17, 18, 21). Here, by directly interrogating the metal identity, redox state, and functional contributions at both the active and allosteric sites, we revise the prevailing model of SAMHD1 and establish a central role for transition metals in its function.

Our findings demonstrate that SAMHD1 is not a magnesium-dependent enzyme (23, 24, 33–38). Instead, it is a transition-metal-dependent hydrolase with Fe^2+^ or Mn^2+^ supporting robust catalysis, and Mg^2+^ alone producing minimal turnover. These findings validate and extend prior structural observations by showing that iron has a central role, not only as a critical component of the catalytic dinuclear core, but also as a potent allosteric activator (17, 18, 21). In this context, SAMHD1 more closely resembles other metallophosphohydrolases in which transition metals, rather than Mg^2+^, drive catalysis (3, 10–13, 15).

A key insight of this work is the role of iron in organizing the dinuclear active site. Spectroscopic and metal reconstitution experiments indicate that iron preferentially occupies one position within the dinuclear center and promotes recruitment of the second metal ion required for catalysis. This apparently cooperative assembly is not equivalently reproduced by manganese, suggesting that iron functions as an architectural element that stabilizes the bimetallic active-site. Similar principles have been observed in other dimetal enzymes, including HD-domain phosphohydrolases and purple acid phosphatases, in which a tightly bound metal ion stabilizes the active-site architecture and promotes binding of a second catalytic metal (10, 15). Within the HD-domain family, fungal SAMHD1 orthologs and the bacterial phosphohydrolase YqeK likewise illustrate that this scaffold can support both homometallic and mixed-metal dinuclear centers to retain catalytic activity (15, 61).

Consistent with this framework, SAMHD1 exhibits substantial plasticity at the second metal-binding site. Heterodinuclear configurations such as Fe-Mg^2+^ and Fe-Mn^2+^ retain catalytic activity, albeit with distinct kinetic properties, indicating that SAMHD1 can function across multiple metalation states. This plasticity suggests that SAMHD1 does not rely on a single rigid cofactor configuration but instead maintains catalytic competence under varying intracellular metal conditions. Such flexibility may be a broader feature of HD-domain enzymes, enabling function across fluctuating metal availability (9, 15, 61).

The catalytic consequences of iron oxidation state in SAMHD1 further depend on the identity of the second metal ion. Diiron centers display pronounced redox sensitivity, with maximal activity observed in the diferrous state, whereas incorporation of a redox-inert metal such as magnesium buffers the enzyme against oxidative inactivation. These findings provide direct experimental evidence that redox state influences SAMHD1 catalysis and suggest that mixed-metal active sites preserve enzymatic function under oxidizing cellular conditions.

In addition to its structural role in the active site, iron also functions as an allosteric activator of SAMHD1 with higher affinity than magnesium. This observation expands the role of transition metals beyond catalysis and suggests that metal availability can influence both assembly and regulation of SAMHD1 activity. Such a property may be particularly relevant in macrophages and other immune cells in which SAMHD1 is highly expressed and both intracellular metal availability and redox conditions fluctuate during infection and immune activation (58, 59, 67–70).

The metal dependence uncovered here also has broader implications for nucleotide metabolism. Mammalian ribonucleotide reductase, which catalyzes de novo dNTP synthesis, is itself a diiron enzyme (39, 41). Our finding that SAMHD1 also relies on iron for efficient dNTP hydrolysis raises the possibility that iron availability could regulate both the synthesis and degradation of deoxyribonucleotides. Although ribonucleotide reductases across biology use diverse metallocofactors (40, 41), the mammalian context suggests an intriguing convergence in which iron contributes to multiple points of dNTP control required for genome stability and antiviral defense.

In summary, our results redefine the metal requirements of SAMHD1 and reveal a metalloregulatory framework for dNTP hydrolysis. Iron emerges as a central determinant of active site assembly, while promiscuity at the second metal-binding site preserves enzymatic function across diverse cellular metal and redox conditions. Beyond its structural role, iron also acts as a high-affinity allosteric activator, linking metal availability to regulation of SAMHD1 activity. This study highlights that transition metals may play a broader role in regulating HD-domain enzymes and the pathways they govern and also reveals a connection between metal homeostasis, nucleotide metabolism, and antiviral defense in SAMHD1 in particular.

## Materials and Methods

### Materials

All chemicals were of high-purity grade and obtained from Fisher Scientific unless otherwise specified.

### Human SAMHD1 Expression

For protein expression, *Escherichia coli* T7 express cells (New England Biolabs) were transformed with a pET-19b plasmid containing the wild-type and variant His_6_-tagged SAMHD1 (Δ112) from *Homo sapiens* and an ampicillin resistance marker (19). The cells were grown in either Luria Bertani (LB) or minimal (M9) media to direct specific metal incorporation. In the case of M9 media the cultures contained 0.5% (w/v) glucose, 90 µM thiamine hydrochloride, 0.1 mM CaCl_2_, 2 mM MgSO_4_, and different transition metal ion salts: 125 μM (NH_4_)_2_Fe(SO_4_)_2_ or 100 μM MnCl_2_, for M9-Fe and M9-Mn respectively. The cells were incubated at 37 °C and 220 rpm until an OD_600_ of 0.6-1.0 was reached. The cultures were cold shocked for 1 h at 4 °C and protein expression was induced by addition of 0.5 mM Isopropyl β-d-1-thiogalactopyranoside (IPTG), and 125 μM (NH_4_)_2_Fe(SO_4_)_2_ or 100 μM MnCl_2_, respectively. After shaking (220 rpm) for 16-20 h at 18 °C, the cells were harvested by centrifugation at 1,600 x g and 4 °C for 20 min, flash frozen in liquid nitrogen, and stored at −80 °C prior to further use.

### Human SAMHD1 Purification

In an O_2_-free glovebox (Coylab), cell pellets were resuspended in lysis buffer (50 mM 4-(2-Hydroxyethyl)piperazine-1-ethanesulfonic acid (HEPES), 300 mM potassium chloride (KCl), 10 mM imidazole, 0.5 mM tris(2-carboxyethyl) phosphine (TCEP), 10% glycerol, pH 7.5) and lysed using a Qsonica sonicator with the addition of 10 mg/L lysozyme, and 0.45 μg/L phenylmethylsulfonyl fluoride (PMSF) prior to cell disruption. Lysed cells were centrifuged at 40,000 x g for 30 min. The clarified lysate was loaded onto a Ni^2+^-NTA immobilized affinity chromatography column (~10 mL resin per 100 mL lysate) equilibrated with the lysis buffer. The column was first washed with lysis buffer and then with wash buffer (50 mM HEPES, 300 mM KCl, 30 mM imidazole, 0.5 mM TCEP, 10% glycerol, pH 7.5). The protein was eluted with elution buffer (50 mM HEPES, 300 mM KCl, 250 mM imidazole, 0.5 mM TCEP, 10% glycerol, pH 7.5). When SAMHD1 was purified in the presence of magnesium (LB-Mg, M9-FeMg) or manganese (M9-FeMn), 4 mM MgCl_2_ or MnCl_2_ respectively were included in the lysis, wash, and elution buffers. The elution was concentrated at 3,900 rpm using a 50 kDa MWCO Amicon Centrifugal Filter Unit (Millipore Sigma, MA). The concentrated protein was loaded onto a HiLoad 16/600 Superdex 200 pg (Cytiva, MA) size exclusion column, which was equilibrated with storage buffer (50 mM HEPES, 300 mM KCl, 0.5 mM TCEP, 10% glycerol, pH 7.5).(19) Fractions containing the pure protein were pooled and further concentrated. Protein purity was estimated by SDS-PAGE with Coomassie staining, and protein concentration was determined using a molar absorption coefficient of 63,260 M^−1^ cm^−1^ at 280 nm (https://web.expasy.org/protparam/) via an Ultraviolet-visible (UV-Vis) spectrophotometer (Agilent, CA). The protein was flash-frozen in liquid nitrogen and stored at −80 °C in air-tight cryovials prior to further use.

### SAMHD1 Chelation

M9-Fe SAMHD1 was reduced with 10 mM sodium dithionite (NaDT) for 10 min in an anaerobic glovebox. Dipicolinic acid (DPA)was then added to a concentration of 1.25 mM, and the protein was left to react at 4 °C overnight. Removal of the NaDT and DPA was achieved under O_2_-free conditions using a NAP-5 desalting column (Cytiva). To ensure optimal chelation, this process was repeated once.

### SAMHD1 Activity Assays

All activity assays were carried out under O_2_-free conditions in an anaerobic glovebox and in the presence of NaDT, unless otherwise specified. SAMHD1 containing storage buffer was preincubated with 10 mM NaDT for 10 min and aliquoted into a 96-well plate. The reactions were initiated by addition of buffer containing the allosteric metal as well as dGTP, the allosteric nucleotide and substrate, to bring the final concentrations in the reaction to 0.5 µM SAMHD1, 0.5 mM NaDT, and 1 mM dGTP, as well as a variable concentration of the allosteric metal (0-5 mM). The reaction was quenched by 1:1 dilution of the reaction with the quench solution containing a 270 µM caffeine internal standard (ε_260_ = 12,399 M^−1^ cm^−1^) and 100 mM potassium hydroxide at 2.5, 5, and 10 min. For the redox-dependent assays, as is SAMHD1 was not preincubated with 10 mM NaDT and oxidized SAMHD1 was preincubated with 7.5 mM hydrogen peroxide, respectively.

### HPLC Analysis

SAMHD1 activity assay samples were analyzed on an Agilent 1260 Infinity II high-performance liquid chromatography system equipped with a 1260 Infinity Photodiode Array Detector WR by monitoring the absorbance at 260 nm (61). The samples were injected on a 150 x 4.6 mm Phenomenex Synergi 4u Fusion-RP 80 Å pore size reverse phase column and run at 60 °C using aqueous (20 mM ammonium acetate pH 4.5, Solvent A) and organic (methanol, Solvent B) mobile phases in a 19 min program consisting of an 11 min gradient from 100% A to 21.5% A and 78.5% B, which ran for a further one min followed by a three min gradient from 21.5% A and 78.5% B to 100% B and a jump to 100% A for the final four min (52). The amount of product (dG) formation was quantified by comparing the integrated absorbances of the dG (ε_260_ = 11,500 M^−1^ cm^−1^) peak to the caffeine (ε_260_ = 12,399 M^−1^ cm^−1^) internal standard peak, and normalizing for their different extinction coefficients at 260 nm.

### Glutaraldehyde Crosslinking

The glutaraldehyde crosslinking was carried out following previously published procedures with some modifications (42). First, 1 µM SAMHD1 was incubated in the crosslinking buffer (50 mM HEPES, 50 mM KCl, pH 7.5) alongside 200 µM dGTP or 1 mM metal, if necessary, for 20 min. The iron condition was performed in an anaerobic glovebox and reduced with 2 mM NaDT to ensure that all the iron was in the ferrous state. Next, 20 µL of 50 mM glutaraldehyde, in water, was added to each sample, and they were incubated at RT for 15 min. The crosslinking reaction was quenched by addition of 8 µL of 1M tris(hydroxymethyl) aminomethane hydrochloride (Tris-HCl) pH 7.5. Finally, 16 µL of 4x Laemmli loading dye was added to each sample, and they were run on Invitrogen NuPAGE 4*-*12%, Bis-Tris, 1.5 mm, 10-well precast gels. The gels were imaged on a ChemiDoc Imager (Bio-Rad).

### Electron Paramagnetic Resonance (EPR)

All samples were prepared in storage buffer (50 mM HEPES, 300 mM KCl, 10% glycerol, 0.5 mM TCEP, pH 7.5) under O_2_-free conditions in an anaerobic glovebox. Reduction with sodium ascorbate was performed with 2 molar eq (with respect to protein concentration) for 10 min at room temperature prior to freezing in liquid nitrogen. EPR spectra were acquired at 10 K on a Bruker E500 Elexsys continuous wave (CW) X-Band spectrometer (operating at approximately 9.36 GHz) equipped with a rectangular resonator (TE102) and a continuous-flow cryostat (Oxford 910) with a temperature controller (Oxford ITC 503).

### EPR Metal Replacement Studies

To monitor metal replacement at the active site of SAMHD1, M9-Fe SAMHD1 was incubated overnight with excess metal salts (100 mM MgCl_2_ 1 mM ZnSO_4_, or 10 mM NiSO_4_). These concentrations were chosen to promote maximal displacement of the second Fe ion while maintaining protein stability. Samples were subsequently frozen in liquid nitrogen and analyzed as described above.

### Inductively Coupled Plasma Atomic Emission Spectrometry (ICP-AES)

Metal analysis of the protein samples was determined by inductively coupled atomic emission spectrometry (ICP-AES) at the Energy and Environmental Sustainability Laboratories (EESL) at the Pennsylvania State University (PA). Samples were prepared by precipitation in 3.5% metal-free nitric acid.

### Ferrozine Assays

Iron stoichiometry was also determined using the colorimetric ferrozine assay. The protein of interest was diluted to 15 μM and precipitated by addition of trichloroacetic acid (Ricca, Tx) to a final concentration of 8% (w/v). After incubation for 20 min, samples were spun at 13,000 x g for 2 min and the supernatant was removed and diluted to a final volume of 500 µL with water. To this sample, 2.3 mM of ascorbic acid, 300 µM of ferrozine, and 18% (v/v) of a saturated ammonium acetate solution were added. The sample was then mixed and the Fe^2+^-ferrozine complex was monitored at 562 nm (ε_562_ = 26,385 M^−1^ cm^−1^).

### ^57^Fe Mössbauer Spectroscopy

Mössbauer samples were prepared from purified M9-Fe SAMHD1 (in storage buffer (50 mM HEPES, 300 mM KCl, 10% glycerol, 0.5 mM TCEP, pH 7.5) under O_2_-free conditions. Samples were reacted with 2 mol eq of sodium ascorbate or 10 mM NaDT for 15 min as indicated. Mössbauer spectra were recorded on a WEB Research (Edina, MN) instrument. The spectrometer used to acquire the weak-field spectra is equipped with a Janis SVT-400 variable-temperature cryostat. The external magnetic field was applied parallel to the γ beam. All isomer shifts are quoted relative to the centroid of the spectrum of α-iron metal at room temperature. Mössbauer spectra were fit using the WMOSS4 Mössbauer Spectral Analysis Software (www.wmoss.org).

## Acknowledgments

The authors thank Dr. Paul Ralifo for providing us access to the EPR spectrometer at the Chemical Instrumentation Center of Boston University (Boston, MA). The authors thank Dr. Laura J. Liermann for ICP-AES analysis at the Laboratory for Isotopes and Metals in the Environment at Pennsylvania State University (University Park, PA). The authors would like to also thank Richard Wang for his assistance and carrying out some of the actitivity assays of the Mn-enriched SAMHD1.

