## Supplemental Information for "Transition metal activation reframes SAMHD1 regulation"

**This PDF file includes:**

Figures S1 to S5

Tables S1 to S5

SI References

Figures

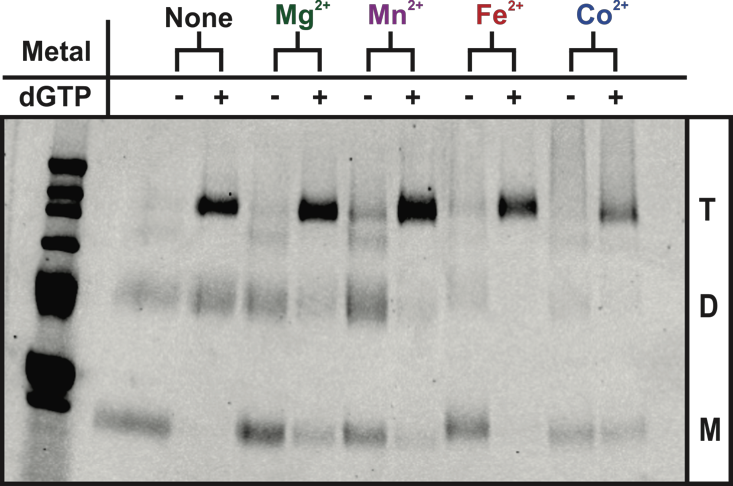

Fig. S1. Glutaraldehyde crosslinking gel monitoring oligomerization of apo WT SAMHD1 by Mg^2+^, Mn^2+^, Fe^2+^, or Co^2+^ with and without dGTP. The T, D, and M labels denote tetramer, dimer, and monomer respectively. Experimental conditions: [apo SAMHD1] = 1 µM, [dGTP] = 200 µM, [Metal] = 1 mM, [NaDT] = 2 mM. Apo-SAMHD1 was generated as described in the materials and methods in the main text.

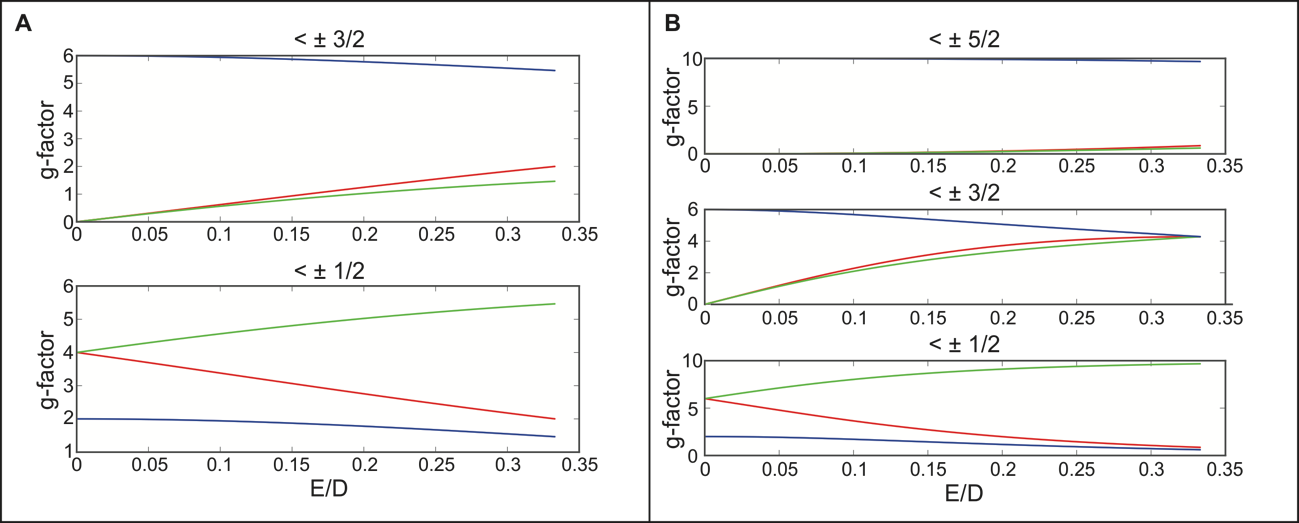

Fig. S2. Rhombograms for spin *S* = 3/2 (A) and *S* = 5/2 (B). The red, green, and blue lines correspond to the x (green), y (red), and z (blue) g-factor components in the three coordinate axes. E and D refer to the rhombic and axial zero field splitting parameters respectively (1).

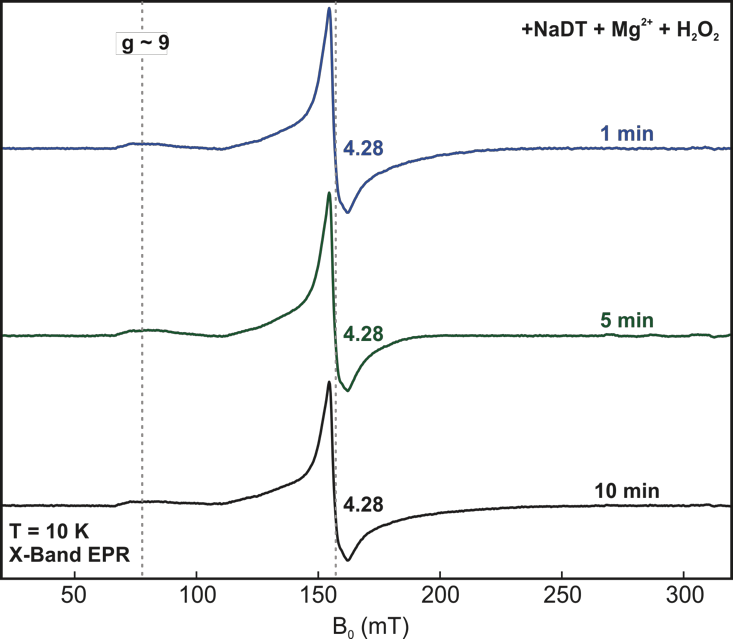

**Fig. S3 EPR spectra monitoring formation of Fe^3+^-Mg^2+^ in M9-Fe WT SAMHD1 by addition of Mg^2+^.** Samples were reduced for one (blue), five (green), or 10 (black) min before addition of 5 mM Mg^2+^ and 1 mM dGTP were added and let react for 10 min, after which hydrogen peroxide was added to accumulate the high-spin Fe^3+^-Mg^2+^ state. Absence of rhombic signals characteristic of the heteronuclear Fe^3+^-Mg^2+^ site confirm that under the activity assay conditions, no detectable replacement of Fe by Mg^2+^ occurs at the active site. Experimental conditions: temperature = 10 K, microwave frequency = 9.38 GHz, microwave power = 2 mW, modulation amplitude = 1 mT. [SAMHD1] = 200 µM. [NaDT] = 1 mM, [Mg] = 5 mM, [H_2_O_2_] = 2 mM for 10 min.

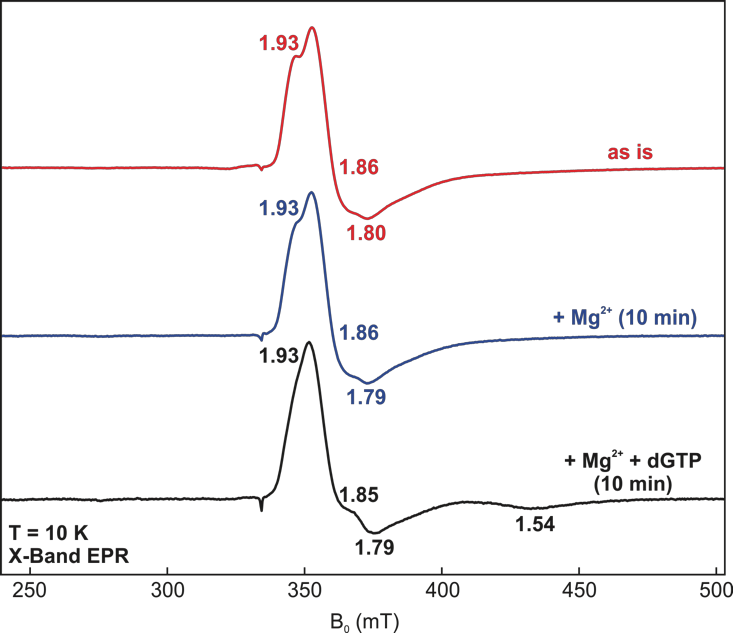

**Fig. S4. EPR spectra of anaerobically isolated M9-Fe WT SAMHD1 showing the effect of Mg²⁺ addition on the Fe^3+^-Fe^2+^ signal.** Spectra are shown for the anaerobically isolated sample (red trace, top), after incubation with 5 mM Mg^2+^ for 10 min prior to freezing (blue trace, middle), and after incubation with 5 mM Mg^2+^ and 1 mM dGTP for 10 min prior to freezing (black trace, bottom). No decrease in the Fe^3+^-Fe^2+^ signal intensity is observed upon Mg^2+^ addition, indicating that under these conditions Mg^2+^ does not replace Fe^2+^ at the active site. In contrast, addition of dGTP results in the appearance of additional resonances consistent with mixed-valent diiron species, suggesting substrate interaction with the active site. Experimental conditions: temperature = 10 K, microwave frequency = 9.38 GHz, microwave power = 2 mW, modulation amplitude = 1 mT, [Mg] = 5 mM, [dGTP] = 2 mM (incubated for 2.5 min prior to freezing).

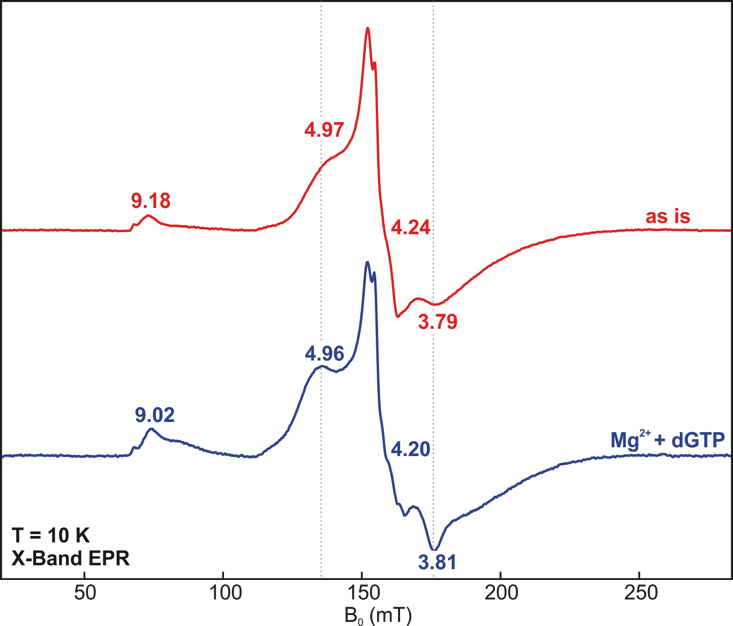

**Fig. S5 Monitoring the effect of dGTP addition to the Fe^3+^-Mg^2+^ signal by EPR spectroscopy.** The high-spin region of the EPR spectrum of the Fe^3+^-Mg^2+^ signal prior to (red trace, top) and after the addition of dGTP (blue trace, bottom). Addition of dGTP leads to changes g-value shifts and changes in rhombicity indicative of dGTP interacting with the Fe^3+^-Mg^2+^ site. Experimental conditions: temperature = 10 K, microwave frequency = 9.38 GHz, microwave power = 2 mW, modulation amplitude = 1 mT. [SAMHD1] = 300 µM, [Mg] = 5 mM, [dGTP] = 10 mM.

Tables

**Table S1. Elemental analysis data (moles of metal per moles of sample) for various SAMHD1 preparations as determined by ICP-AES or the ferrozine assay.** LOD indicates the measurement was below the limit of detection of the ICP-AES instrument. Units are moles of metal per mole of protein or nucleotide (for the dGTP control).

| **Sample** | **Ca** | **Co** | **Cu** | **Fe** | **Mg** | **Mn** | **Ni** | **Zn** |
| --- | --- | --- | --- | --- | --- | --- | --- | --- |
| WT (apo)  (ferrozine) |  |  |  | 0.09 |  |  |  |  |
| WT (M9-Fe) | 6.44 | <0.017 (LOD) | <0.016 (LOD) | 1.12 | 0.28 | <0.018 (LOD) | 0.72 | 0.16 |
| WT (LB) | 2.21 | <0.017 (LOD) | <0.016 (LOD) | 0.04 | 0.19 | <0.018 (LOD) | 0.05 | 0.61 |
| WT (LB- -Mg) | 2.58 | <0.017 (LOD) | <0.016 (LOD) | 0.11 | N/A | 0.05 | 0.52 | 0.84 |
| H233A (M9-Fe) | 2.15 | <0.017 (LOD) | <0.016 (LOD) | 0.26 | 0.18 | 0.03 | 0.16 | 0.58 |
| WT (M9-Mn) | 4.79 | <0.017 (LOD) | <0.016 (LOD) | 0.18 | 0.33 | 0.77 | 0.25 | 0.10 |
| H233A (M9-Mn) | 5.07 | <0.017 (LOD) | <0.016 (LOD) | 0.15 | 0.37 | 0.71 | 0.04 | 0.16 |
| WT (M9-FeMg)* | 2.84 | <0.017 (LOD) | <0.016 (LOD) | 0.52 | N/A | 0.027 | 0.80 | 0.26 |
| WT (M9-FeMn) | 2.07 | <0.017 (LOD) | <0.016 (LOD) | 0.40 | 0.08 | 0.53 | 1.93 | 0.35 |
| C522S (M9-Fe) | 4.17 | <0.017 (LOD) | <0.016 (LOD) | 0.30 | 0.28 | <0.018 (LOD) | 0.31 | 0.99 |
| C522S (LBMg added)*  (ferrozine) |  |  |  | 0.328 |  |  |  |  |
| dGTP Control | 0.085 | <0.0002 (LOD) | <0.0002 (LOD) | <0.0002 (LOD) | 0.006 | <0.0002 (LOD) | <0.0002 (LOD) | 0.002 |

^* LBMg and FeMg refer to sample preparations in which magnesium was added in the purification buffer as described in the materials and methods and therefore the amount of magnesium is omitted as it does not reflect magnesium bound to SAMHD1.^

**Table S2. Observed dGTP hydrolysis rates for wild-type and H233A Mn-enriched SAMHD1.** Reactions were performed at room temperature and hydrolysis rates were determined from time-course measurements collected over a 10 min interval. Experimental conditions: [M9-Mn SAMHD1] = 0.5 µM, [dGTP] = 1 mM, [Mn] = 2 mM or [Mg] = 2.5 mM. Results are reported as averages ± the standard error of the mean (n = 4).

| **Condition** | **k_obs_ (s^-1^)** |
| --- | --- |
| WT + Mn | 2.13 ± 0.5 |
| WT + Mg | 1.86 ± 0.6 |
| H233A + Mn | 0.02 ± 0.03 |
| H233A + Mg | n.d. |

**Table S3. SAMHD1 catalytic parameters for dGTP hydrolysis for different metal configurations of the active and allosteric sites.** Experimental conditions: [SAMHD1] = 0.5 µM, [dGTP] = 1 mM, [NaDT] = 0.5 mM. [Mg^2+^] was titrated from 0-2.5 mM, while [Fe^2+^] and [Mn^2+^] were titrated from 0-1 mM. Results are reported as averages ± the standard error of the mean (n = 3).

| **Condition** | **k_obs_ (s^-1^)** | **K_act_ (µM)** |
| --- | --- | --- |
| Fe^2+^-X^2+^ active site / Mg^2+^ allosteric activator | | |
| Fe^2+^-Mg^2+^ | 1.80 ±0.2 | 257 ± 80 |
| Fe^2+^-Fe^2+^ | 1.82 ± 0.2 | 277 ± 80 |
| Fe^2+^-Mn^2+^ | 0.77 ± 0.08 | 417 ± 120 |
| Fe^2+^-Mg^2+^ active site / X^2+^ allosteric activator | | |
| Mg^2+^ | 1.80 ± 0.2 | 257 ± 80 |
| Fe^2+^ | 1.42 ± 0.05 | 68 ± 10 |
| Mn^2+^ | 1.42 ± 0.07 | 47 ± 10 |

**Table S4. Distribution of diiron redox states in M9-Fe SAMHD1 as determined by ^57^Fe Mössbauer spectroscopy.** Mössbauer parameters are quoted at T = 80 K. [ascorbate] = 2 mol. eq with respect to protein, [NaDT] = 10 mM.

| **Sample** | **Site** | **δ (mm/s)** | **∆E_Q_ (mm/s)** | **Area (%)** |
| --- | --- | --- | --- | --- |
| as isolated | | | | |
| Fe^3+^-Fe^3+^ | Fe^3+^ | 0.53 | 0.90 | 49 |
| Fe^3+^-Fe^2+^ | Fe^3+^ | 1.17 | 2.6 | 15 |
|  | Fe^2+^ | 0.56 | 0.89 | 15 |
| + ascorbate | | | | |
| Fe^3+^-Fe^3+^ | Fe^3+^ | 0.53 | 0.90 | 10 |
| Fe^3+^-Fe^2+^ | Fe^3+^ | 1.18 | 2.54 | 28 |
|  | Fe^2+^ | 0.56 | 0.89 | 28 |
| Fe^2+^-Fe^2+^ | Fe^2+^ | 1.25 | 2.74 | 30 |
| + NaDT | | | | |
| Fe^2+^-Fe^2+^ | Fe^2+^ | 1.25 | 2.74 | 100 |

**Table S5. Observed dGTP hydrolysis rates for the C522S SAMHD1 enriched in diiron and Fe-X active site configurations and under different redox conditions.** Experimental conditions: [SAMHD1] = 0.5 µM, [dGTP] = 1 mM, [Mg] = 5 mM, [H_2_O_2_] = 7.5 mM, [NaDT] = 10 mM. M9-Fe indicates SAMHD1 grown in M9-media with supplemental iron as described in the materials and methods. LB-Mg indicates SAMHD1 grown in LB and purified in the presence of magnesium as described in the materials in methods. Results are reported as averages ± the standard error of the mean (M9-Fe oxidized, as is, n = 5), (M9-Fe reduced, n = 6), (LB-Mg oxidized, as is, n = 7), (LB-Mg reduced, n = 8).

| **Condition** | **k_obs_ (s^-1^)** |
| --- | --- |
| M9-Fe  (H_2_O_2_ oxidized) | 0.24 ± 0.03 |
| M9-Fe | 0.31 ± 0.03 |
| M9-Fe  (NaDT reduced) | 0.62 ± 0.08 |
| LB-Mg  (H_2_O_2_ oxidized) | 1.12 ± 0.1 |
| LB-Mg | 1.21 ± 0.1 |
| LB-Mg  (NaDT reduced) | 1.51 ± 0.1 |
